# Hidden Structural Bias in Proteomics: Sonication-induced Selective Fragmentation of Intrinsically Disordered Regions

**DOI:** 10.64898/2026.07.14.738389

**Authors:** Megumi Narita, Tatsuya Yamakawa, Rika Nishimura, Mio Iwasaki

**Affiliations:** Center for iPS Cell Research and Application, Kyoto University, Kyoto 606-8507, Japan

**Keywords:** Sonication, Intrinsically Disordered Regions (IDRs), Protein Fragmentation, Structural Bias, PEPPI-MS, Sample Preparation, Quantitative Proteomics

## Abstract

Sonication is a fundamental technique in proteome sample preparation, primarily used for protein solubilization and shearing of genomic DNA. Although the mechanical shearing of DNA is well-characterized, its unintended impact on protein structural integrity remains a significant “blind spot” in high-throughput analytical workflows. In this study, we systematically investigated sonication-induced protein fragmentation by combining gel-based fractionation (PEPPI-MS) with sequence-level compositional analysis and bioinformatic mapping.

Our results demonstrate that sonication does not significantly alter overall proteome identification or the recovery of membrane proteins; however, it induces extensive and non-random protein fragmentation. Sonication caused an approximately three-fold increase in the abundance of >45 kDa protein-derived fragments migrating into the <40 kDa fraction, and 1,620 high-molecular-weight (MW) proteins were uniquely detected in the lower-MW fraction upon sonication, an eight-fold increase over non-sonicated controls. Peptide-level amino acid composition analysis revealed subtle but directional shifts in the sonication-derived fragments.

This residue-level signature is reinforced by two orthogonal structural analyses (MobiDB peptide-level mapping and protein-level profiling using metapredict V3 software), which show that sonication-susceptible proteins harbor more than twice the disordered content of length-matched controls (median 40% vs. 18%).

This study identifies a previously unrecognized “structural bias” whereby intrinsically disordered region (IDR)-rich proteins are selectively compromised during sample preparation. Because these fragments are indistinguishable from enzymatic digestion products in conventional bottom-up proteomics, the underlying structural damage is effectively masked in global quantitative datasets, potentially distorting biological interpretations related to protein size, isoforms, and stability, particularly for IDR-rich classes, such as transcription factors and signaling molecules. We propose that optimizing and standardizing sonication parameters is essential for ensuring the accuracy and reproducibility of quantitative proteomic analyses.

## INTRODUCTION

Proteomics has become an indispensable tool for large-scale protein expression profiling, with recent advancements in mass spectrometry achieving sensitivities that enable single-cell analyses (1). However, as analytical sensitivity increases, the impact of sample preparation on quantitative accuracy becomes more pronounced, making the optimization of these workflows a critical step (2). Among the current methodologies, sonication—using a Bioruptor or probe-type system—is a standard procedure in proteome sample preparation. A Europe PMC survey indicates that sonication is described in the Methods section of approximately 22% of all mass spectrometry-based proteomics articles published since 2000 (Table S9), underscoring its status as a de facto standard step in modern proteome sample preparation. It is primarily employed to enhance protein solubilization through structural denaturation and to shear residual genomic DNA, thereby reducing sample viscosity. Reflecting the growing recognition of sonication as a key parameter in proteome workflows, the Clinical Proteomic Tumor Analysis Consortium (CPTAC) recently incorporated a sonication step into its tissue lysis protocol and demonstrated improved detection of membrane-bound and DNA-binding proteins (3).

Sonication is also intentionally used in genomic studies, such as Chromatin Immunoprecipitation (ChIP), to fragment DNA (4, 5), where consistent shearing requires careful parameter optimization, with sonication itself acknowledged as a potential pitfall when conditions are not controlled (6). Beyond these intended applications, however, the impact of sonication on the structural integrity of proteins remains poorly characterized. Earlier biophysical work demonstrated that sonication can induce amyloid-like aggregation across structurally diverse proteins via acoustic cavitation (7), and a more recent report indicated that myofibrillar proteins can be fragmented under combined sonication and heat treatment (8). To our knowledge, no proteome-wide study has systematically evaluated whether sonication selectively damages specific protein classes during sample preparation. Consequently, the extent to which this phenomenon occurs across the entire proteome and the resulting systematic biases it may introduce into high-throughput proteome datasets remain significant blind spots.

In this study, we investigated whether sonication induces significant protein cleavage by employing a gel-based fractionation approach (PEPPI-MS), which uniquely enables direct visualization of molecular weight shifts at the protein level prior to enzymatic digestion. Although structural mass spectrometry methods are increasingly used to characterize IDRs and intrinsically disordered proteins (IDPs) (9), the extent to which standard sample preparation steps can selectively damage these flexible regions has not been investigated. To dissect the structural specificity of sonication-induced cleavage, we combined this gel-based readout with two complementary sequence-level analyses: (i) compositional analysis of the amino acid frequencies of peptides identified across conditions, and (ii) mapping of identified peptides onto predicted disordered regions retrieved from MobiDB. Our findings demonstrate that sonication selectively cleaves proteins within their intrinsically disordered regions (IDRs), introducing a previously unrecognized “structural bias” into quantitative proteome datasets.

## Experimental Procedures

### Experimental Design and Statistical Rationale

Two lysate conditions were compared throughout this study: sonicated and non-sonicated, each prepared from the human induced pluripotent stem cell (iPSC) line, 692D2. For each condition, proteins were fractionated by SDS-PAGE into a Gel Top (>40 kDa) and a Gel Bottom (<40 kDa) fraction, with each fraction separately analyzed by LC-MS/MS, yielding four sample groups (sonicated Gel Top, sonicated Gel Bottom, non-sonicated Gel Top, non-sonicated Gel Bottom). Three independent biological replicates were prepared for each condition from separate cultures of the same iPSC line harvested on different days (replicate batches 250528, 250610, and 250625), yielding a total of 12 LC-MS/MS runs (2 conditions × 2 fractions × 3 replicates). The non-sonicated samples served as the baseline control for sonication-specific effects. This replicate number (n = 3 biological replicates per condition) was chosen as the minimum required to estimate within-condition variance and to enable two-condition statistical testing while remaining practical for the gel-based fractionation. Per-residue and physicochemical-class amino acid frequencies (Figure 3) were compared between conditions within each gel fraction using two-sample Student’s t-tests (n = 3 vs. n = 3) with Benjamini–Hochberg false-discovery-rate correction across the 20 amino acids within each fraction, and effect sizes were quantified as Cohen’s d. The number of peptides mapped to IDRs (Figure 4B, C) was compared using log10-transformed Welch’s two-tailed t-tests (n = 3). For protein-level disorder metrics (Figure 2I–K), continuous metrics were compared between the two groups using two-sided Mann– Whitney U tests, with Cliff’s delta as the effect size. The proportion of proteins containing long IDR was compared using Fisher’s exact test. P-values from these comparisons were adjusted using the Benjamini–Hochberg procedure. To exclude protein length as a confounder, the sequence length distributions of the two groups were compared using the Mann–Whitney U test. The remaining comparisons in Figure 1 used two-tailed Student’s t-tests (n = 3). Statistical analyses were performed using Python (NumPy and SciPy).

**Figure 1.**
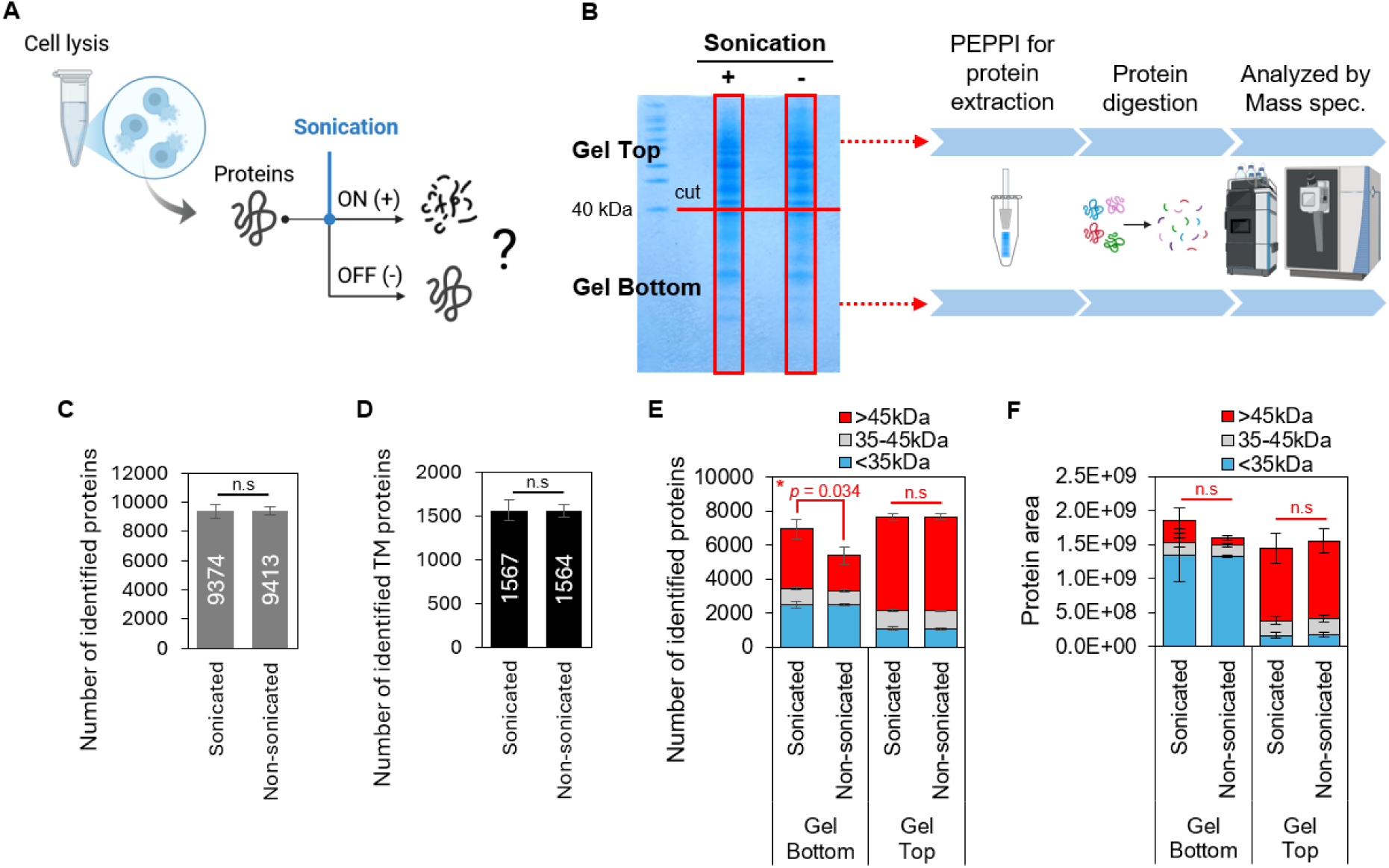
Sonication-induced protein fragmentation in cell lysates. (A) Experimental design for comparing the proteomic profiles of sonicated and non-sonicated cell lysates. Human iPSCs were lysed in PTS buffer and heat-shocked (95°C, 5 min) to inactivate endogenous proteases, followed by sonication (Bioruptor, 10 cycles) or maintenance on ice (non-sonicated). (B) Schematic representation of the PEPPI-MS workflow. Proteins were separated by SDS-PAGE, and segments above 40 kDa (Gel Top) and below 40 kDa (Gel Bottom) were excised for proteomic analysis. (C) Total number of identified proteins merged from Gel Top and Gel Bottom fractions (n=3). (D) Identification of transmembrane proteins under both conditions. Transmembrane proteins were defined by searching the Swiss-Prot database for human entries annotated as transmembrane. (E) Number of identified proteins in Gel Bottom and Gel Top fractions, categorized by their database-registered molecular weights (>45, 35–45, and <35 kDa; the 35–45 kDa band represents the ambiguous boundary zone around the 40 kDa gel cut). (F) Total quantitative abundance of proteins in Gel Bottom and Gel Top fractions, grouped by molecular weights. Statistical significance was assessed using a two-tailed Student’s t-test. P-values shown in red indicate comparisons of proteins with database-registered molecular weights >45 kDa between sonicated and non-sonicated samples (n.s., not significant; *p < 0.05). Data are presented as mean ± SD (n=3).

**Figure 2.**
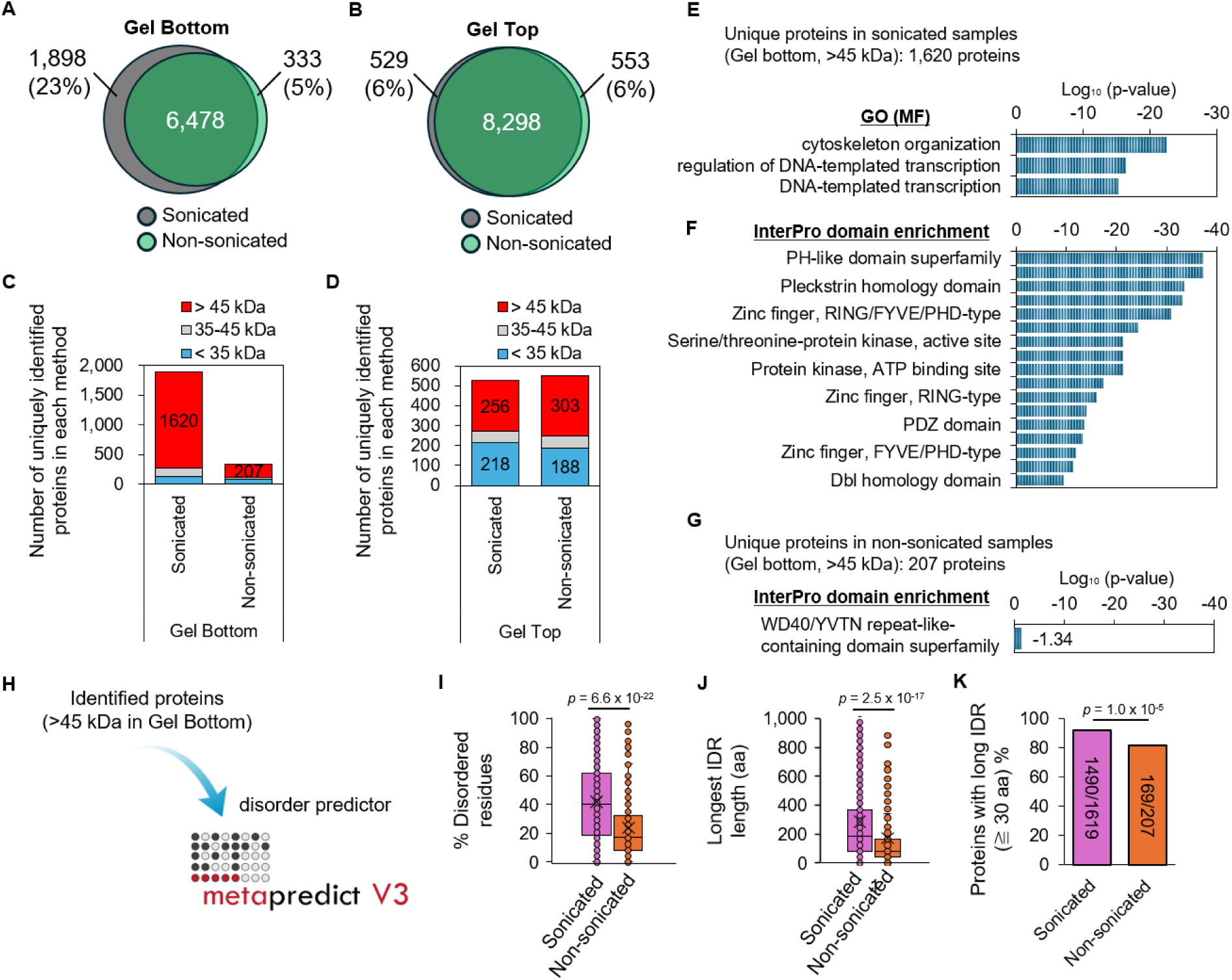
Characterization of structurally susceptible proteins during sonication. (A, B) Venn diagrams showing the overlap of identified proteins between sonicated and non-sonicated samples in Gel Bottom (A) and Gel Top (B) fractions. (C, D) Number of uniquely identified proteins categorized by molecular weight in Gel Bottom (C) and Gel Top (D) fractions. (E) Gene Ontology (GO) Molecular Function (MF) enrichment analysis of the 1,620 high-molecular-weight proteins (>45 kDa) uniquely identified in the sonicated Gel Bottom fraction. (F, G) InterPro domain enrichment analysis of uniquely identified proteins in the sonicated (F) and non-sonicated (G) Gel Bottom fractions. (H) Schematic of the protein-level disorder analysis: full-length sequences of proteins uniquely identified in the Gel Bottom fraction (>45 kDa) under each condition were scored for intrinsic disorder using metapredict V3 software. (I–K) Per-protein disorder metrics for proteins uniquely identified in the sonicated (n = 1,590) versus non-sonicated (n = 206) Gel Bottom (>45 kDa) fractions. (I) Percentage of disordered residues per protein (metapredict score > 0.5; median 40.2% vs. 17.5%). (J) Length of the longest contiguous disordered segment per protein (amino acids; median 185 vs. 79 aa). (K) Proportion of proteins containing at least one long intrinsically disordered region (IDR ≥ 30 aa); numbers on the bars indicate proteins meeting the criterion / total proteins (1,461/1,590 vs. 168/206; 91.9% vs. 81.6%, respectively). In (I, J), box plots show the median (center line), mean (×), and interquartile range (box), with individual proteins overlaid as points; magenta, sonicated; white, non-sonicated. Statistics: (I, J) two-sided Mann–Whitney U test with Cliff’s delta as the effect size; (K) Fisher’s exact test; P-values were corrected using the Benjamini–Hochberg procedure. Counts in (I–K) reflect unique UniProt entries after de-duplication of the upstream protein lists; see Methods and Table S4.

**Figure 3.**
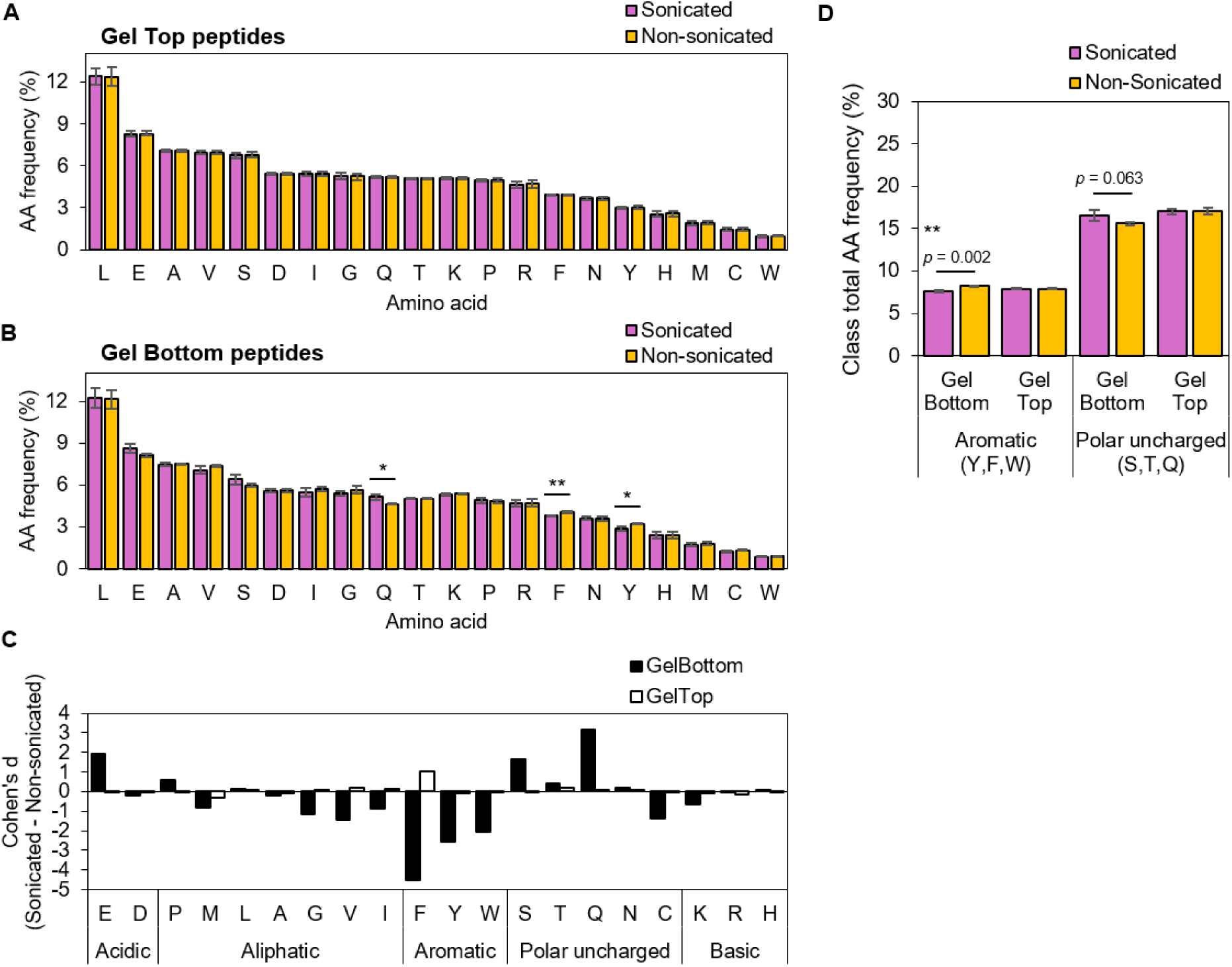
Sonication-derived peptides exhibited subtle compositional shifts consistent with IDR enrichment. All panels in this figure show the identified peptides (peptide-level analysis). Per-residue amino acid frequencies were computed for peptides uniquely identified in either the sonicated or non-sonicated samples within each fraction (n = 3 replicates per condition). (A) Mean per-residue amino acid frequency for peptides uniquely identified in the Gel Top fraction (>40 kDa) of sonicated and non-sonicated samples. (B) Mean per-residue amino acid frequency for peptides uniquely identified in the Gel Bottom fraction (<40 kDa). (C) Effect-size summary across all 20 amino acids shows that Cohen’s d values are large for Gel Bottom (yellow) but essentially zero for Gel Top (black), with consistent directionality (depletion of order-promoting residues and enrichment of disorder-promoting residues). (D) Pooled class comparison demonstrates the IDR-like signature: aromatic class (Y, F, W) is significantly depleted (Cohen’s d = −5.85; p = 0.002) and polar uncharged class (S, T, Q) shows an enrichment trend (Cohen’s d = +2.08; p = 0.063) in sonication-derived peptides from Gel Bottom fractions, while neither class shows any change in Gel Top control fractions. Data are shown as mean ± SD (n = 3). Significance markers (Student’s t-test): ** p < 0.01; * p < 0.05.

**Figure 4.**
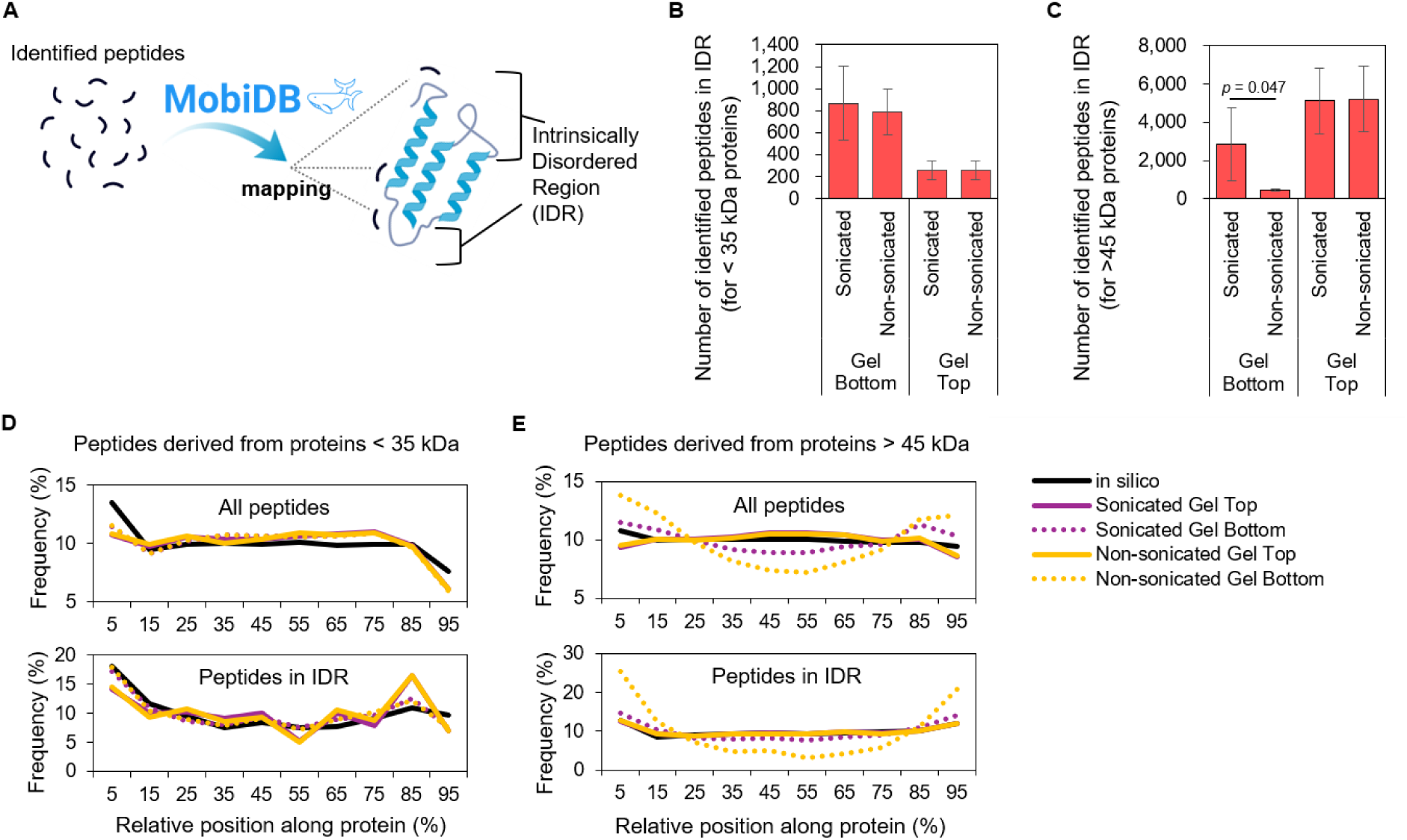
Preferential cleavage of intrinsically disordered regions (IDRs) by sonication. All panels in this figure show the analysis of identified peptides (peptide-level analysis). (A) Bioinformatic workflow for mapping identified peptides to MobiDB disorder annotations. (B, C) Number of identified peptides mapped within MobiDB-annotated IDRs in Gel Bottom and Gel Top fractions for low-molecular-weight (<35 kDa) proteins (B) and high-molecular-weight (>45 kDa) proteins (C). The Gel Top fraction served as a control and showed comparable counts between conditions. Data are presented as mean ± standard deviation (SD). (D, E) Relative positional distribution of identified peptides for proteins <35 kDa (D) and >45 kDa (E). For each panel, the upper subpanel shows all identified peptides, and the lower subpanel shows the subset of peptides mapped to MobiDB-lite-annotated IDRs. Error bars are omitted for clarity. Across three biological replicates, the standard deviation per bin did not exceed 1.2% for all-peptide and 3.0% for IDR analyses. The per-bin mean and SD values are provided in Table S7.

### Cell culture

Human iPSCs (692D2) were maintained on culture plates coated with iMatrix-511 silk (Nippi) in StemFit AK02N medium (Ajinomoto). Cell passaging was performed as previously described (10). Briefly, cells were washed with Dulbecco’s Phosphate Buffered Saline (D-PBS; Nacalai Tesque) and incubated with TrypLE Express (Thermo Fisher Scientific) at 37°C for 10–15 min. Cells were then dissociated into single cells by pipetting and collected in Dulbecco’s Modified Eagle Medium/Ham’s F-12 (DMEM/F-12; Wako) supplemented with 0.1% bovine serum albumin (BSA; Wako). After centrifugation and removal of the supernatant, cells were resuspended in StemFit AK02N medium. Following cell counting, cells were seeded onto new plates at an optimal density in the presence of 0.5 µg/mL iMatrix-511 (Nippi) and 10 µM Y-27632 (Nacalai Tesque).

### Protein Extraction and PEPPI-MS Analysis

Cells were washed once with ice-cold PBS (Nacalai Tesque) and lysed directly with ice-cold PTS buffer (12 mM sodium deoxycholate, 12 mM sodium lauroyl sarcosinate, and 100 mM Tris-HCl, pH 9.0) supplemented with 1% phosphatase and protease inhibitors (Sigma-Aldrich). Lysates were collected by scraping into pre-chilled 1.5 mL Protein LoBind tubes (Eppendorf). Immediately after collection, lysates were heated at 95°C for 5 min to ensure complete protein denaturation and inactivation of endogenous proteases.

Following heat treatment, samples were divided into two groups: sonicated and non-sonicated groups. For the sonicated group, lysates were sonicated using a Bioruptor (Sonicbio, #UCW-310) with a cycle of 30 s ON / 30 s OFF for 10 cycles at “High” power while maintaining the temperature at 4°C. The non-sonicated group was maintained on ice for the same duration. Protein concentrations were determined using the BCA protein assay (Thermo Fisher Scientific).

Protein extraction from polyacrylamide gels was performed using the previously described PEPPI method (11, 12). Briefly, 5 µg of protein samples were separated on 4-12% Bis-Tris gradient gels (1 mm thick, NuPAGE, Thermo Fisher Scientific) using NuPAGE MES running buffer. Molecular weight markers were HiMark Unstained Protein Standard (Thermo Fisher Scientific, #LC5688) loaded in a parallel lane to define the 40 kDa excision position. Following electrophoresis, gel regions corresponding to proteins <40 kDa and >40 kDa were excised and treated as Gel Top and Gel Bottom samples downstream. After Coomassie Brilliant Blue staining with EzStain Aqua (ATTO), gel pieces were crushed, homogenized, and filtered through a 3-kDa molecular weight cut-off (MWCO) centrifugal filter (Amicon Ultra, #UFC5003). Recovered intact proteins were used for subsequent analyses.

### Sample Preparation for Proteomic Analysis

Sample preparation was performed as previously described (13), with modifications to the protein aggregation capture (PAC) method (14, 15). Briefly, hydrophilic and hydrophobic Sera-Mag SpeedBead carboxylate-modified magnetic particles were mixed in equal proportions, washed three times with distilled water, and reconstituted at a concentration of 15 µg/µL. Protein samples were reduced with 20 mM tris(2-carboxyethyl)phosphine (TCEP) and alkylated with 30 mM iodoacetamide (IAA). Alkylated proteins were then mixed with 20 µL of reconstituted beads in the presence of 80% (v/v) ethyl alcohol to induce aggregation. After two washes with 80% ethyl alcohol, the supernatant was removed, and the beads were resuspended in 100 µL of 50 mM Tris-HCl (pH 8.0). Proteolysis was performed by adding 2 µL of Trypsin/Lys-C Mix and incubating at 37°C overnight. The resulting peptides were acidified with 20 µL of 5% trifluoroacetic acid (TFA) and desalted using in-house SDB-XC StageTips (16). Finally, peptide concentrations were determined using a quantitative fluorometric peptide assay (Thermo Fisher Scientific).

### Proteomic Analysis

For the timsTOF Pro 2 system (Bruker), peptide samples were loaded and separated using a nanoElute (Bruker) and an Aurora column (25 cm length, 75 μm i.d., Ion-Opticks, Melbourne, Australia). The mobile phase, comprising 0.1% formic acid (solution A) and 0.1% formic acid in acetonitrile (solution B), was delivered at a flow rate of 400 nL/min. For the 120-min gradient, solution B was increased from 2% to 17% over 60 min, 17% to 25% over 30 min, 25% to 37% over 10 min, and then to 80% over 10 min, followed by an additional 10 min at 80% for washing the column. Columns were washed with 80% solution B after analysis. The applied spray voltage was 1,400–1,500 V, and the interface heater temperature was 180°C. For data-independent acquisition (DIA), the Parallel Accumulation–Serial Fragmentation (PASEF) method (diaPASEF) was used (17). For the diaPASEF settings, a cycle time of 1.74 s with 16 diaPASEF scans was employed, with an MS/MS isolation width of 28 m/z, precursor ion ranges of 391–1,175 m/z, and ion mobility ranges of 0.69–1.39 V·s/cm². No iRT or other retention-time standards (e.g., spike-in peptides) were used; retention-time alignment was performed empirically by DIA-NN. Samples were not randomized for acquisition order; they were injected in the order of replicate batches and conditions as listed in Supplemental Table S8. Each biological replicate was analyzed by a single LC-MS injection (no technical replicates); analytical reliability was therefore assessed by 1% precursor- and protein-group-level FDR filtering, by match-between-runs–based consistency of MaxLFQ quantification across the twelve files, and by the reproducibility of quantitative values across the three biological replicates.

DIA data were processed with DIA-NN v1.9 (18) operated in its library-prediction (“library-free”) mode (--fasta-search with --gen-spec-lib --predictor). A predicted in silico spectral library was first generated from the human UniProt/Swiss-Prot database (release 2024_02; 42,911 entries comprising 20,795 canonical Swiss-Prot human proteins, 22,093 isoforms, 23 TrEMBL entries, and 381 common contaminants) (19) using DIA-NN’s built-in deep-learning models for fragment intensity, retention time, and ion mobility predictions. Library generation parameters were as follows: trypsin specificity with up to one missed cleavage, N-terminal methionine excision, peptide length 7–30 residues, precursor m/z 300–1,800, precursor charge 1–4, fragment ion m/z 200–1,800, cysteine carbamidomethylation as a fixed modification, and methionine oxidation and protein N-terminal acetylation as variable modifications (up to one variable modification per peptide). The initial predicted library contained 5,907,665 target precursors corresponding to 20,638 unique proteins (isoforms grouped at the protein level by DIA-NN).

Decoy precursors were generated using DIA-NN’s default reverse-sequence strategy. DIA spectra were searched against this predicted library with peptidoform scoring enabled (-- peptidoforms), heuristic protein grouping (--relaxed-prot-inf), retention-time profiling across runs (--rt-profiling), and DIA-NN’s cross-run “reanalyse” (match-between-runs) mode (--reanalyse). Precursor and fragment mass tolerances were determined automatically per run by DIA-NN. A first pass identified high-confidence precursors that were used to build an empirically aligned spectral library, which was then used in a second pass for final identification and quantification. Identifications were filtered at 1% false-discovery rate at the precursor level (q ≤ 0.01) and at 1% global FDR at the protein-group level, yielding approximately 9,200 protein groups per replicate batch (8,912–9,471 across three biological replicate batches). Retention-time and ion-mobility tolerances were likewise auto-optimised by DIA-NN at each run. Extracted-ion chromatograms for single-peptide protein identifications can be reproduced from the deposited DIA-NN report files and the raw .d data at jPOSTrepo (JPST004418) / ProteomeXchange (PXD076301) using DIA-NN with the parameters specified above.

### Amino acid composition analysis of identified peptides

Peptides that could not be assigned to a single protein (i.e., mapping to more than one UniProt accession) were excluded from the analysis. Peptide sequences were stripped of all modifications, and only the 20 standard amino acid residues were retained for compositional calculations. Within each MS run, peptides were considered “detected” when their intensity was greater than zero, and redundant entries sharing an identical stripped amino acid sequence (e.g., different charge states or modification variants of the same peptide) were collapsed to a single occurrence so that each unique peptide sequence contributed once per run.

Peptides were analyzed separately for the two gel regions: the upper region (Gel Top), corresponding to proteins of approximately >40 kDa, and the lower region (Gel Bottom), corresponding to proteins of approximately <40 kDa. For each peptide, the relative frequency of each amino acid was calculated as the residue count divided by the peptide length and expressed as a percentage. For each MS run, the amino acid frequency was obtained as the mean across all unique detected peptides (each peptide was weighted equally, irrespective of length). This yielded three independent replicate values for each amino acid and gel region for the sonicated and non-sonicated samples.

For each amino acid, frequencies were compared between sonicated and non-sonicated samples within each gel region using a two-sample Student’s *t*-test (equal variance assumed, n = 3 versus n = 3). To account for multiple tests, *p*-values were adjusted within each gel region (20 amino acids) using the Benjamini–Hochberg false discovery rate procedure. In parallel, residues were grouped into physicochemical classes, and the class-total frequency (aromatic, Y + F + W; polar uncharged, S + T + Q) was computed per run and compared between conditions in the same manner. Effect sizes were quantified as Cohen’s *d* (sonicated minus non-sonicated, using the pooled standard deviation) for each amino acid in each gel region. Data are presented as mean ± standard deviation (SD) of three replicates; significance is indicated as \**p* < 0.05, \*\**p* < 0.01. All calculations were performed using Python (NumPy and SciPy).

### Peptide-level mapping to intrinsically disordered regions (IDRs)

To investigate the structural distribution of identified peptides, human protein entries (Proteome ID: UP000005640) were retrieved from the MobiDB database (version: mobidb_search_2025-07-09T13-22-33-HUMAN). For comparison, a reference set of *in silico* digested peptides was generated from the UniProt/Swiss-Prot human proteome (release 2024_02, including isoforms and contaminants (19)) using the following criteria: trypsin-like cleavage at Lys/Arg residues (excluding the RP motif) and mass range of 600–4000 Da. This analysis was performed at the peptide level. Each identified peptide was mapped against the predicted disordered regions defined by the “pred.dis.mobi.dblite” dataset in MobiDB, and the relative frequency and spatial distribution of these peptides within disordered sequences were calculated to characterize their localization patterns.

To compare the number of IDR-mapped peptides between sonicated and non-sonicated samples (Figure 4B, C), peptide counts were log10-transformed prior to statistical analysis. Log transformation was applied to address the non-normal distribution and unequal variance characteristics of MS-derived count data, consistent with standard practices in quantitative proteomics. Differences between groups were assessed using Welch’s two-tailed t-test (n = 3 replicates per condition), which does not assume equal variance between groups.

### Protein-level disorder profiling using metapredict

To assess intrinsic disorder at the protein level, per-residue disorder scores were predicted using metapredict V3 software (20). The analyzed set consisted of all proteins uniquely identified in either the sonicated or non-sonicated Gel Bottom fraction whose UniProt-annotated molecular weight exceeded 45 kDa, corresponding to the size class shown to be selectively fragmented by sonication (Figure 2C; n = 1,590 sonicated, n = 206 non-sonicated). For each protein, three features were calculated from its full-length sequence: (i) the fraction of residues predicted to be disordered using a threshold score of 0.5 (frac_idr), (ii) the length of the longest contiguous disordered segment (longest_idr), and (iii) the number of long disordered segments ≥ 30 amino acids (n_long_idr). Group-level distributions of these continuous metrics were compared using two-sided Mann–Whitney U tests, and Cliff’s delta was computed as a nonparametric effect size. The proportion of proteins containing at least one long IDR was assessed using Fisher’s exact test. P-values for the four comparisons were corrected using the Benjamini–Hochberg procedure. To control for sequence-length confounding, sequence length distributions of the two groups were compared using the Mann–Whitney U test.

## RESULTS & DISCUSSION

### Sonication induces the fragmentation of high-molecular-weight proteins

To investigate the impact of sonication on protein integrity, freshly cultured human iPSCs (692D2) were lysed and subjected to two experimental conditions: sonicated and non-sonicated (Figure 1A). To rule out enzymatic degradation as a confounder, lysates from both conditions were heat-inactivated (95°C, 5 min) immediately after lysis and prior to sonication, ensuring that any observed fragmentation reflects mechanical shearing rather than residual protease activity. Proteins were separated by molecular weight using SDS-PAGE, and the gel was excised at the 40 kDa mark. Proteins were then extracted from segments above 40 kDa (Gel Top) and below 40 kDa (Gel Bottom) using the PEPPI method, followed by comprehensive proteomic analysis (Figure 1B).

The total number of identified proteins, calculated by merging the Gel Top and Gel Bottom data for each replicate, showed no significant difference between sonicated and non-sonicated samples (Figure 1C, Table S1). As sonication is commonly used to enhance protein solubilization, we also compared the recoverability of transmembrane proteins. However, no significant difference was observed between the two conditions (Figure 1D, Table S2). These observations suggest that sonication does not alter the overall accessibility of the proteome in conventional bottom-up proteomics workflows.

Next, we compared the number of identified proteins and their quantitative values based on their registered molecular weights. Because manual excision of the gel at the 40 kDa marker and protein migration in SDS-PAGE are not perfectly precise, a ±5 kDa buffer margin was used.

Proteins were categorized by their database-registered molecular weight into >45 kDa, 35–45 kDa, and <35 kDa groups (Figure 1E, Table S3), with the 35–45 kDa band treated as an ambiguous boundary. A significant difference was detected in the number of proteins >45 kDa identified in the Gel Bottom fraction (sonicated: 3,492 proteins; non-sonicated: 2,084 proteins). Quantitative analysis revealed that the total abundance of >45 kDa proteins in Gel Bottom fractions was approximately three-fold higher in the sonicated samples than in the non-sonicated controls (sonicated: 3×10⁸; non-sonicated: 1×10⁸; Figure 1F, Table S3). In contrast, Gel Top fractions showed negligible differences in both identification and quantification between the two conditions. Additionally, no significant differences were observed for proteins <35 kDa or within the 35–45 kDa range in the Gel Bottom fraction (Figure 1E, F, Table S3). These results indicate that while sonication does not alter overall proteome identification efficiency, it induces substantial protein fragmentation, specifically causing high-molecular-weight proteins to fragment and migrate into lower-molecular-weight fractions. These observations establish the analytical framework used throughout the remainder of this study: the Gel Top fraction, dominated by intact high-molecular-weight proteins, serves as an internal negative control, whereas the Gel Bottom fraction harbors the sonication-derived fragments characterized in the subsequent analyses.

### Selective fragmentation of high-molecular-weight proteins reveals structural susceptibility

To further characterize the impact of sonication on the proteome, we analyzed the overlap of identified proteins between sonicated and non-sonicated samples (Figure 2A, B). In the Gel Top fraction, only approximately 6% of proteins were uniquely identified under either condition (Figure 2B). In contrast, the Gel Bottom fraction from the sonicated samples contained 1,898 unique proteins (23% of the total), a significantly higher proportion than that in the non-sonicated control (Figure 2A). When categorized by molecular weight, the number of proteins >45 kDa uniquely identified in the Gel Bottom fraction was approximately eight-fold higher with sonication (1,620 proteins) than without sonication (207 proteins) (Figure 2C). In contrast, no such difference was observed in the Gel Top fraction across all molecular weight ranges (Figure 2D). These data confirm that sonication partially cleaves at least 1,620 high-molecular-weight proteins, releasing fragments that migrate into the lower-molecular-weight range without affecting the identification of the intact proteome in the Gel Top fraction.

To identify features of these sonication-susceptible proteins, we performed functional enrichment analysis on the 1,620 proteins (>45 kDa) uniquely identified in Gel Bottom fractions of sonicated samples (Figure 2E, F). Gene Ontology molecular function (GO-MF) analysis revealed an enrichment of proteins associated with the cytoskeleton and DNA-dependent RNA transcription (Figure 2E). Furthermore, InterPro domain analysis showed that proteins containing specific structural motifs, such as the PH-like domain superfamily and zinc fingers, were significantly enriched (Figure 2F). Notably, both domain families are frequently embedded within or flanked by extended IDRs in transcription factors and signaling molecules. In contrast, the non-sonicated control (207 proteins) showed no significant GO enrichment and was characterized primarily by WD40/YVTN repeat-containing domains (Figure 2G), which adopt rigid β-propeller architectures with minimal disorder, consistent with these proteins representing endogenous degradation products rather than sonication-induced fragments.

To directly assess the structural nature of these sonication-susceptible proteins, we profiled intrinsic disorder across their full-length sequences using metapredict V3 software, an orthogonal deep learning-based predictor (Figure 2H–K, Table S4). This protein-level analysis was restricted to proteins with molecular weights > 45 kDa, matching the size class shown to be selectively fragmented by sonication (Figure 2C). Proteins uniquely identified in the sonicated Gel Bottom (n = 1,590 proteins) contained more than twice the disorder content of those uniquely identified in the non-sonicated control (n = 206 proteins) (median frac_idr 40.2% vs. 17.5%; Figure 2I). Sonication-susceptible proteins also harbored substantially longer maximum disordered segments (median 185 vs. 79 amino acids; Figure 2J) and were significantly more likely to contain at least one long IDR ≥ 30 residues (91.9% vs. 81.6%; Figure 2K). Critically, the two protein groups had essentially identical sequence-length distributions (median 779 vs. 772 aa, p = 0.61), excluding the well-known correlation between protein length and disorder content as a confounding factor. Together, these domain- and disorder-level features of the sonication-susceptible proteins led us to hypothesize that sonication preferentially targets disordered protein segments. To test this hypothesis, we performed two complementary peptide-level analyses: a residue-composition analysis (Figure 3) that asks whether sonication-derived peptides display the canonical compositional signature of IDRs without invoking any structural annotation and a structural-mapping analysis (Figure 4) that directly assigns each identified peptide to MobiDB-annotated disordered regions of its source protein. The convergence of these annotation-free and annotation-based tests, together with the protein-level disorder analysis already presented above, establishes IDR enrichment as a robust feature of sonication-induced cleavage.

### Sonication-derived peptides exhibit a subtle but directional compositional shift consistent with IDR enrichment

To characterize sonication-induced fragmentation at the residue level, we computed the amino acid composition of peptides uniquely identified in the sonicated or non-sonicated samples of both Gel Top and Gel Bottom fractions (using the Gel Top fraction as the internal negative control defined above). Consistent with this expectation, none of the 20 amino acids exhibited statistically significant compositional differences between the conditions in the Gel Top fraction (Figure 3A, Table S5), confirming that sonication does not perturb the residue-level composition of structurally intact proteins.

In contrast, the Gel Bottom fraction, enriched in sonication-derived fragments, exhibited subtle but directionally consistent shifts (Figure 3B, Table S5). The order-promoting aromatic residue Phe (F) was the most strongly affected, being depleted in the sonication-derived peptides (p = 0.005). The aromatic residue Tyr (Y, depleted) and disorder-promoting polar uncharged residue Gln (Q, enriched) also reached nominal significance (p < 0.05). Although no residue survived the Benjamini–Hochberg correction at n = 3 (minimum q = 0.10), the effect sizes were exceptionally large (e.g., Phe: Cohen’s d = −4.52; Gln: +3.18; Tyr: −2.58), and all 20 residues directionally mirrored the canonical IDR compositional signature (Figure 3C, Table S5). To capture this signal at the level of biochemical residue classes, we pooled aromatic (Y, F, and W; canonical order-promoting) and polar uncharged (S, T, and Q; canonical disorder-promoting) residues. In the Gel Bottom fraction, the aromatic class was significantly depleted in sonication-derived peptides (p = 0.002), whereas the polar uncharged class showed a corresponding enrichment trend (Figure 3D, Table S5). No such shifts were observed in the Gel Top fraction (both classes |d| < 0.15; p > 0.88), confirming that the residue-level signature is specific to sonication-derived fragments rather than to overall sample handling. These compositional shifts provide annotation-free sequence-level evidence that sonication-induced fragments preferentially originate from IDR-rich segments. To complement this with a structurally explicit test, we next mapped each identified peptide onto MobiDB-annotated disordered regions of its source protein.

### Sonication-induced cleavage preferentially targets intrinsically disordered regions (IDRs)

To elucidate the structural distribution of sonication-induced fragments at the peptide level, we mapped the identified peptides to disorder annotations retrieved from the MobiDB database (Figure 4A, Table S6). Quantitative peptide-level analysis revealed that the high-molecular-weight (>45 kDa) proteins migrating to the Gel Bottom fraction under sonication contained 6.3-fold more IDR-mapped peptides than those in the non-sonicated control (mean 2,811 vs. 445; p = 0.047, log10-transformed Welch’s t-test; Figure 4C). In contrast, no such enrichment was observed for proteins <35 kDa, where peptide counts were nearly identical between conditions (mean 827 vs. 752; p = 0.83; Figure 4B), establishing that the IDR enrichment is specific to the size class of proteins selectively fragmented by sonication.

Next, we analyzed the positional distribution of the identified peptides along the protein sequences, considering all identified peptides and the subset mapped to MobiDB-annotated IDRs (Figure 4D, E, Table S7). For proteins <35 kDa, the identification patterns largely matched the *in silico* digestion model and remained consistent across sonication and gel fractions in both all-peptide and IDR-restricted analyses (Figure 4D). However, a striking deviation was observed for proteins >45 kDa. While the patterns were consistent across both Gel Top fractions and the sonicated Gel Bottom fraction, the non-sonicated Gel Bottom fraction—representing basal, endogenous protein fragments—showed a uniquely N/C-terminus-enriched distribution that was reproduced when restricting the analysis to IDR-mapped peptides (Figure 4E). In this control, peptides were predominantly detected at the N- or C-termini, consistent with N/C-terminal proteolytic processing or co-translational degradation observed in steady-state proteomes. In contrast, sonication-induced fragments were characterized by a high frequency of peptides originating from internal regions of the proteins, and this internal redistribution was even more pronounced in the IDR-restricted analysis, further supporting the IDR specificity of sonication-induced cleavage. This selective cleavage pattern parallels observations from limited proteolysis mass spectrometry (LiP-MS), in which enzymatic cleavage sites preferentially localize to the flexible and disordered regions of proteins (21), reflecting the inherent accessibility of IDRs to external cleavage agents, whether enzymatic or mechanical. Together, residue composition (Figure 3), peptide-level IDR enrichment and positional localization (Figure 4B–E), and protein-level disorder profiling (Figure 2H–K) provide concordant evidence that sonication does not randomly degrade proteins but specifically targets the structurally flexible IDRs within high-molecular-weight proteins.

## LIMITATIONS

This study has several limitations. The analysis was performed on a single cell type (human iPSCs) using a single lysis buffer (PTS) and a single sonication device (Bioruptor). Whether the same patterns occur with probe-type sonicators or in other biological systems remains to be determined. While our conclusions on IDR selectivity are supported by two independent disorder-prediction frameworks (MobiDB consensus peptide-level annotations and metapredict deep learning-based protein-level predictions), further cross-validation using additional predictors, such as IUPred or PONDR, would help solidify the structural interpretation. Future work extending this analysis across diverse sample preparation conditions and alternative mechanical disruption techniques will help generalize the implications of the structural bias described here.

## CONCLUSIONS

In summary, we report that sonication, a near-ubiquitous step in proteomic sample preparation, induces non-random, structurally selective fragmentation of high-molecular-weight proteins.

Three independent and convergent analyses (peptide-level amino acid composition, peptide-level MobiDB mapping, and protein-level metapredict profiling) established that the cleavage preferentially targets intrinsically disordered regions (IDRs). Our findings extend the isolated ChIP-context observation by Pchelintsev et al. (6) into a systematic, proteome-wide phenomenon and identify IDRs as the structural locus of cleavage, providing a mechanistic explanation for size dependence. Because sonication-derived fragments are indistinguishable from enzymatic peptides in standard bottom-up workflows, this phenomenon constitutes a previously unrecognized “structural bias” that may systematically distort quantitative comparisons of IDR-rich proteins, including many transcription factors and signaling molecules. Therefore, we advocate for the careful optimization and reporting of sonication parameters, and the recognition of the trade-off between detection depth and structural preservation, to safeguard the reproducibility and biological interpretation of mass spectrometry-based proteomics.

## AUTHOR CONTRIBUTIONS

Megumi Narita: Investigation, Methodology. Tatsuya Yamakawa: Formal analysis. Rika Nishimura: Formal analysis. Mio Iwasaki: Conceptualization, Formal analysis, Funding acquisition, Project administration, Supervision, Writing – original draft, Writing – review & editing. All authors have read and agreed to the published version of this manuscript.

## ACKNOWLEDGMENT

We thank members of the Mio Iwasaki laboratory at CiRA (Kyoto University) for fruitful discussions, Kelvin Hui for critical reading of the manuscript, Mayu Terakawa for making the graphical abstract, and S. Aimi, A. Matsuzaki, M. Matsui, and S. Takeshima for administrative support. Illustrations were created using BioRender.

## DECLARATION OF GENERATIVE AI AND AI-ASSISTED TECHNOLOGIES IN THE MANUSCRIPT PREPARATION PROCESS

During the preparation of this manuscript, the authors used an AI assistant (Anthropic Claude) to assist with data analysis scripting and language editing. All AI-assisted outputs and codes were reviewed, verified, and edited by the authors, who took full responsibility for the content of the published article.

## DATA AND MATERIALS AVAILABILITY

All data supporting the findings of this study are available in the article and its supplementary materials. The raw mass spectrometry proteomics data and search results were deposited in the ProteomeXchange Consortium via the jPOSTrepo partner repository(22) (https://repository.jpostdb.org/) with the dataset identifier JPST004418 (PXD076301). For peer review, reviewer access credentials were provided during manuscript submission. A file mapping each raw data file to its condition, gel fraction, and replicate, together with the software and versions used, is provided in Supplemental Table S8. Source data represented in the figures are provided in Supplemental Tables (Figure 1: S1–S3; Figure 2: S1 and S4; Figure 3: S5; Figure 4: S6, S7). All per-peptide identification scores (Q.Value, Global.Q.Value, CScore), modifications, precursor charges, and per-protein accession lists are available in the DIA-NN report files deposited at jPOSTrepo (JPST004418) / ProteomeXchange (PXD076301). Precursor m/z values, the number of distinct peptides per protein, and sequence coverage can be derived from these files and the UniProt Swiss-Prot human reference database used for searching (release and entry counts specified in Methods).

## ABBREVIATIONS

IDR: intrinsically disordered region
IDP: intrinsically disordered protein
PEPPI-MS: Passively Eluting Proteins from Polyacrylamide gels as Intact species for mass spectrometry
LC-MS/MS: liquid chromatography-tandem mass spectrometry
DIA: data-independent acquisition
diaPASEF: data-independent acquisition parallel accumulation-serial fragmentation
FDR: false discovery rate
BH: Benjamini-Hochberg
iPSC: induced pluripotent stem cell
PTS: phase-transfer surfactant
BCA: bicinchoninic acid
SD: standard deviation

## SUPPORTING INFORMATION

Table S1. The list of uniquely identified proteins shown in Figure 1C, 1E–F, and 2A–D.

Table S2. The list of uniquely identified transmembrane proteins shown in Figure 1D.

Table S3. The original number of uniquely identified proteins and quantified areas shown in Figure 1E-F.

Table S4. The per-protein disorder metrics calculated by metapredict V3 software for proteins uniquely identified in sonicated and non-sonicated Gel Bottom (>45 kDa) fractions, corresponding to Figure 2H–K. After de-duplication, n = 1,590 (sonicated) and n = 206 (non-sonicated) unique proteins were analyzed.

Table S5. Amino acid frequency data of uniquely identified peptides shown in Figure 3, including per-replicate frequencies, t-test p-values, BH-corrected q-values, and Cohen’s d effect sizes for both Gel Top and Gel Bottom fractions.

Table S6. The list of uniquely identified peptides shown in Figure 4B–E.

Table S7. The per-bin positional distribution of identified peptides corresponding to Figure 4D, E. For each protein molecular-weight class (<35 kDa and >45 kDa), each gel fraction (Gel Top and Gel Bottom), each condition (sonicated and non-sonicated), and both peptide subsets (all identified peptides and the subset mapped to MobiDB-lite-annotated IDRs), the table reports the mean frequency (%), the standard deviation across three biological replicates, and the per-replicate values for each 10%-wide positional bin (N- to C-terminus) along the source protein sequence.

Table S8. Mapping of each raw mass spectrometry data file to its condition, gel fraction, and replicate, with the corresponding processed and result files and software versions.

Table S9. Europe PMC survey of sonication usage in the proteomics methods sections.

